# FATE (Fish Aquarium with a Turbulent Environment): a turbulence-control facility for quantifying fish–flow interactions and collective behavior

**DOI:** 10.64898/2026.03.25.714166

**Authors:** Michael A. Calicchia, Rui Ni

**Affiliations:** Department of Mechanical Engineering, The Johns Hopkins University, 3400 N. Charles Street, 21218, MD, USA

**Keywords:** Turbulent Flows, Jet Array, Collective Behavior, Fish Locomotion

## Abstract

Despite its ubiquity in natural flows, the effects of turbulence on fish locomotion and behavior remain poorly understood. The prevailing hypothesis is that these effects depend on the spatial and temporal scales of the turbulence relative to the fish’s size and swimming speed. But in conventional facilities, turbulence usually increases with mean flow, which forces higher swimming speeds and can leave these relative scales unchanged. We therefore present a novel experimental facility that leverages a jet array to decouple the turbulence from the mean flow and systematically control its scales. This approach allows the ratio of turbulent to fish inertial scales to be varied over an order of magnitude, providing a controlled framework for quantifying fish–turbulence interactions. The facility also supports experiments probing strategies fish may use to cope with turbulence, including collective behaviors. Insights from this work have broader implications for ecological studies and engineering applications, including the design of effective fishways and bio-inspired underwater vehicles.

## Introduction

Understanding how fishes sense and interact with the surrounding flow has long attracted interest, motivated by both fundamental questions in animal locomotion and by applications in the design of fishways (Silva et al., 2011, 2012a,b; Knapp et al., 2019; Syms et al., 2021), hydrokinetic turbines (Courtney et al., 2022; Müller et al., 2023; Brown, 2023; Macias et al., 2024), and bio-inspired underwater robots (Jiang et al., 2019; Li et al., 2022b; Othman et al., 2023; Koiri et al., 2025). Recent work has demonstrated that flow conditions strongly influence behavior, as fish exploit flow heterogeneity to reduce energy expenditure (Di Santo and Goerig, 2025). To save energy, fish seek out reduced velocity regions (Webb, 2006; Johansen et al., 2007; Liao, 2007; Webb and Cotel, 2010) or coherent vortical structures, such as Kármán vortex streets behind bluff bodies (Liao et al., 2003a,b; Beal et al., 2006; Taguchi and Liao, 2011; Liao and Akanyeti, 2017) or neighboring fish (Li et al., 2020; Thandiackal and Lauder, 2023; Yang and Zeng, 2025). Conversely, fish commonly avoid regions with elevated turbulence (Smith et al., 2005, 2006; Cotel et al., 2006; Goettel et al., 2015; Smith et al., 2014; Hockley et al., 2014; Tan et al., 2019; Li et al., 2022a; Liu et al., 2023), likely because its multiscale and stochastic nature can significantly increase the fish’s cost of transport (Hinch and Rand, 1998; Pavlov et al., 2000; Enders et al., 2003; Roche et al., 2014; Maia et al., 2015; Agbeti et al., 2024; Zhang et al., 2024). Beyond increasing energetic cost, turbulence can also have broader ecological consequences, influencing predator–prey interactions (Higham et al., 2015; Ishikawa et al., 2025), prolonging migration times (Hinch and Rand, 1998; Silva et al., 2011, 2012a,b; Duguay et al., 2018; Knapp et al., 2019), and, in extreme cases, causing bodily injury or mortality (Căda et al., 1999; Odeh et al., 2002; Neitzel et al., 2004).

However, not all studies have reported negative effects of turbulence on swimming performance (Nikora et al., 2003; Ogilvy and Dubois, 1981; van der Hoop et al., 2018), raising the question of when does turbulence actually negatively impact fish locomotion. The answer is expected to depend on the flow scales relative to fish size and swimming speed, yet the relevant nondimensional quantities governing fish–turbulence interactions remain uncertain.

Early work suggested that fish are most affected when the integral length scale is comparable to their body length (Lupandin, 2005). Eddies much smaller than the body are thought to act as minor perturbations that fish can easily resist, while eddies much larger behave more like a slowly varying mean flow, producing little destabilizing effects (Webb and Cotel, 2010; Cotel and Webb, 2015). But, as the eddies approach roughly two-thirds of the body length, their ability to destabilize the fish becomes more significant. (Lupandin, 2005; Tritico and Cotel, 2010; Silva et al., 2012a; Cotel and Webb, 2012). More recent studies have demonstrated that length-scale comparisons alone are insufficient to determine when turbulence will negatively affect the fish. Instead, they proposed comparing the linear and angular momentum, orientation, and periodicity of the eddies relative to that of the fish and its motion (Cotel and Webb, 2012, 2015; Webb and Cotel, 2010; Lacey et al., 2011; Trinci et al., 2017).

These studies primarily focus on the extreme conditions under which turbulence causes fish to lose postural control and be advected downstream, which is known as spilling. Although limited data is available, this framework does appear to adequately predict when turbulence will cause spills. However, this is a situation fish would actively avoid in natural settings. Thus, these studies may not fully capture the conditions under which fish begin to actively avoid turbulent regions during routine navigation and may not fully explain when turbulence begins to negatively affect the fish’s swimming performance.

For example, in Zhang et al. (2024), giant danio (*Devario aequipinnatus*) were exposed to turbulence generated by a passive grid across a range of swimming speeds. The turbulence at all speeds was characterized by a fixed integral length scale of roughly 1.5 cm (20-30% of the fish body length) and nearly constant turbulence intensity (*I* ∼ 20%). Thus, estimates of the linear and angular momentum of the eddies relative to that of the fish yielded ratios well below unity and remained approximately constant for all swimming speeds. Based on these commonly invoked scaling arguments, the turbulence would therefore be expected to have minimal impact on swimming performance with little variation across speeds. Contrary to this expectation, the total cost of transport increased significantly at higher speeds, and post-exercise recovery time was substantially longer after swimming in turbulent versus laminar flows (Zhang et al., 2024). These observations indicate that the conventional metrics fail to explain both when turbulence begins to affect the fish and why its impact becomes progressively stronger at higher swimming speeds, highlighting the need for alternative descriptors of the fish-turbulence interactions.

The results suggest that it may be the coupling between the mean flow and the turbulence characteristics that governs a fish’s navigation strategy and energetic costs. However, most prior experiments have been limited in scope: turbulence fluctuations were typically coupled to the mean flow because they were generated by flow over obstacles such as bricks (Smith et al., 2005, 2006), arrays of cylinders (Tritico and Cotel, 2010), passive grids (Zhang et al., 2024; Ishikawa et al., 2025), or orifices (Silva et al., 2011, 2012a,b). Consequently, increasing turbulence amplitude in these setups also required higher swimming speeds, which in turn increased the momentum of the fish. Ideally, the fish’s momentum should remain constant while the turbulent eddies are systematically varied until their momentum becomes comparable to that of the fish. Addressing this limitation requires experiments in which turbulence characteristics can be independently controlled from the mean flow, enabling systematic investigation of how turbulence alone influences locomotion and behavior.

This could be achieved by utilizing methods that inject momentum into the flow instead of siphoning energy from the mean flow to drive the turbulence. Such methods include the use of jets/jet array (Gad-El-Hak and Corrsin, 1974; Odeh et al., 2002; Bellani and Variano, 2014; Carter et al., 2016; Masuk et al., 2019; Harding et al., 2019; Tan et al., 2023), oscillating grids (De Silva and Fernando, 1994; Villermaux et al., 1995; Cheng and Law, 2001; Stiansen and Sundby, 2001), and propellers/paddles (Zimmermann et al., 2010; Maia et al., 2015). More recently, a jet array has been utilized to study fish behavior in high turbulence levels at relatively low mean flows (Harding et al., 2019). However, a systematic change of the turbulence level was not performed.

To improve our understanding of how turbulence affects fish, we introduce a novel experimental facility that can independently control the turbulence intensity from the mean flow enabling a systematic investigation of both their coupled and individual effects on fish locomotion and behavior. This capability provides a controlled framework for identifying the mechanisms that make swimming in turbulence energetically costly and thereby promote avoidance. Furthermore, the facility can also be used to study another another equally-important aspect of fish-turbulence interaction: understanding the strategies fish employ to avoid turbulent regions or to mitigate their effects when avoidance is not possible. A potential mechanism that fish may utilize to navigate turbulence is collective behavior. Although the ecological and evolutionary benefits of schooling have been extensively studied, including enhanced mating success (Pavlov and Kasumyan, 2000), improved foraging efficiency (Abrahams and Colgan, 1985), and reduced predation risk (Pitcher et al., 1982), very few studies have considered collective swimming as a strategy for coping with environmental turbulence.

Two primary benefits of collective behavior are especially relevant in this context: (1) reduced energy expenditure (Weihs, 1973; Daghooghi and Borazjani, 2015; Marras et al., 2015; Nadler et al., 2016; Ashraf et al., 2017; Li et al., 2019; Schumann et al., 2023; Zhang and Lauder, 2023; Pan et al., 2024; Menzer et al., 2025; Zhou et al., 2025b) and (2) enhanced navigation and migration efficiency in noisy environments (Berdahl et al., 2013; Granger and Johnsen, 2022). To date, however, only a limited number of experiments have directly examined live fish schools navigating turbulent flows. These studies indicate that the presence of conspecifics can modulate fish trajectories (Goettel et al., 2015), improve avoidance of unfavorable flow conditions (Li et al., 2023), and significantly reduce the cost of transport (Zhang et al., 2024). Nevertheless, the mechanisms by which social and hydrodynamic cues interact to shape navigation strategies or unlock significant energy savings in turbulence remain largely unresolved.

Therefore, there exists a need to advance our understanding of how fish perceive, interpret, and respond to turbulent flow environments, both individually and collectively. Specifically, it remains necessary to elucidate the mechanisms that make swimming in turbulence energetically costly, to determine how mean flow and turbulence jointly influence decision-making, and to identify the strategies individual and groups of fish employ to cope with complex hydrodynamic conditions. In this paper, we discuss how the new experimental facility can help achieve these goals and present avenues for future research directions.

### FATE: Fish Aquarium with a Turbulent Environment

Understanding fish-turbulence interactions requires experimental capabilities that allow independent control of the mean flow and turbulence characteristics. To this end, we developed a new facility designed to systematically investigate fish behavioral responses to well-controlled turbulent conditions. As shown in Fig. 1, the facility, termed FATE (**F**ish **A**quarium with a **T**urbulent **E**nvironment), is an open, recirculating water channel, comprising of a main flow loop, flow conditioning section, turbulence generator and controller, and integrated filtration and heating systems. The structure is constructed of non-corrosive materials, including fiberglass, acrylic, and Polyvinyl chloride (PVC) and circulates reverse osmosis (RO) water, which is treated and heated to 26^°^C to maintain satisfactory water quality for the fish. Three 1200W heaters placed in the inlet and exit sections are used to maintain the water temperature.

**Figure 1.**
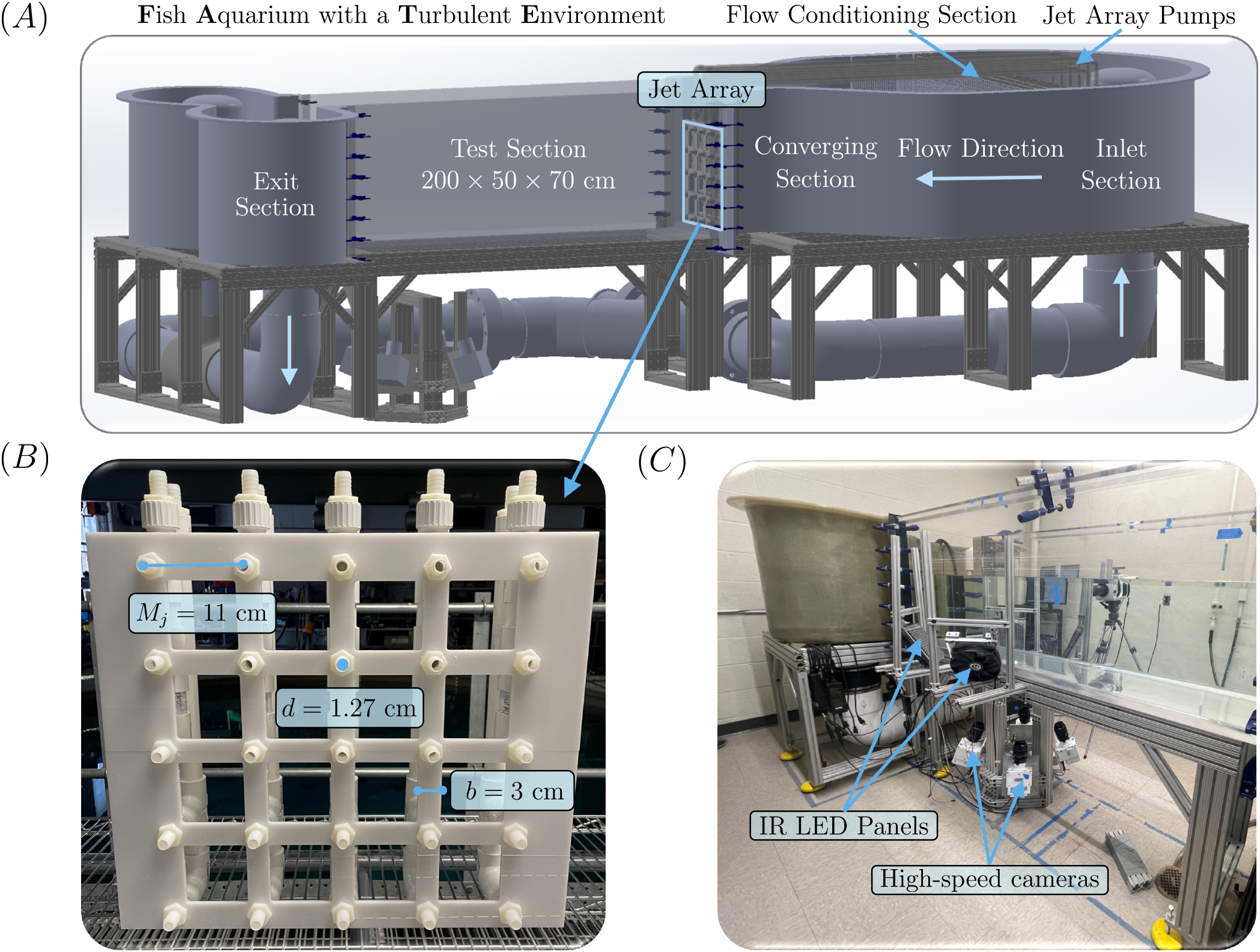
(A) 3D model of FATE (**F**ish **A**quarium with a **T**urbulent **E**nvironment) highlighting all of the major features. (B) Picture of the jet array installed in the facility demonstrating the jet spacing, nozzle diameter, and bar width of the passive grid. (C) Picture of FATE depicting the high-speed cameras and infrared (IR) LED panels used to quantify the turbulence via 3D particle tracking.

Flow is driven by an axial pump with a 23 cm diameter propeller powered by a 1 hp motor, producing mean velocities in the test section of ⟨*u*_1_⟩ = 0.05−0.25 ms^−1^, depending on the pump frequency setting and water level. The acrylic test section measures 200 × 50 × 70 cm providing sufficient downstream distance for turbulence development and a large volume for fish to swim freely with minimal boundary effects.

#### Flow conditioning section

In order to systematically control the turbulence within the tank, the background turbulence inherent to the system must first be minimized. To achieve this, a honeycomb followed by two fine-mesh screens were installed upstream of the test section to suppress large-scale eddies and reduce overall turbulence levels. The cell size of the honeycomb *M*_*h*_ is recommended to be smaller than the smallest lateral wavelength of the velocity variation, estimated to be roughly 1/150 of the settling chamber diameter (Barlow et al., 1999; Mehta and Bradshaw, 1979). Accordingly, a value of 1.27 cm was selected. The length-to-cell-size ratio *L*_*h*_*/M*_*h*_ is another important design parameter, as too large a value may generate excessive pressure losses. Furthermore, the laminar jets exiting the honeycomb can become unstable, generating additional unwanted turbulence with increasing length (Loehrke and Nagib, 1976). An optimal ratio in the range 5–10 is recommended (Barlow et al., 1999; Mehta and Bradshaw, 1979), and thus a honeycomb length *L*_*h*_ = 8.48 cm was chosen.

To further attenuate turbulence introduced by instabilities in these laminar jets, a screen with a mesh size of approximately *M*_*h*_*/*4 was placed roughly 10*M*_*h*_ downstream of the honeycomb (Loehrke and Nagib, 1976). Finally, a second screen with solidity *σ* = 35% and a Reynolds number based on the wire diameter *Re*_*d*_ = 13.3 at the maximum flow speed was installed approximately 15 cm upstream of the contraction section (Barlow et al., 1999; Mehta and Bradshaw, 1979). The flow conditioning section provides a baseline turbulence intensity of approximately 1–3% in the test section, depending on the mean flow rate.

#### Turbulence generator and controller: jet array

Turbulence is generated using a jet array composed of a passive grid, with a solidity *σ* = 60%, into which an array of nozzles is embedded. The nozzles are spaced by *M*_*j*_ = 11 cm and have diameter *d* = 1.27 cm, giving a spacing-to-diameter ratio of *M*_*j*_*/d* = 8.66. The spacing ensures full coverage of the channel cross-section while limiting the total number of jets and associated power requirements, and the chosen spacing-to-diameter ratio is consistent with values used in comparable jet array studies (Gad-El-Hak and Corrsin, 1974; Masuk et al., 2019; Tan et al., 2023). The jet array installed in the facility can be seen in Fig. 1(B).

The passive grid was fabricated by laser cutting an acrylic sheet, while the jet array was assembled from 1.25 in. PVC piping, flexible tubing, and barbed fittings, resulting in a cost-efficient and modular design. Ten independent jet lines are each driven by a dedicated 95 W aquarium pump located in the tank inlet, with each pump supplying two nozzles. The top row of nozzles is blocked, yielding 20 continuously operating jets during experiments. Each pump provides 20 discrete power settings to regulate flow rate, and inline ball valves allow fine adjustment of the jet velocity. By attaching flow meters to each nozzle, the maximum variation in speed between any two jets was measured to be roughly 2.8%, which is comparable to that reported in Gad-El-Hak and Corrsin (1974). The current system can achieve jet speeds ranging from 0.5 to 2.75 ms^−1^.

#### Flow measurements

To quantify the turbulence, the water is seeded with polyamide tracer particles for performing Lagrangian particle tracking. The particles have a nominal radius *a* of 50 *µ*m with a density close to 1.1 g*/*cm^3^. The particle response time *τ*_*p*_ [*τ*_*p*_ = 2*a*^2^Δ*ρ/*(9*νρ*_*f*_)] is approximately 0.56 ms. The Kolomogorov time scale 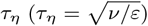 is approximately 95 ms. The Stokes number (*St* = *τ*_*p*_*/τ*_*η*_) is the ratio of these two timescales, which is much smaller than unity, indicating that these particles can safely be treated as tracers.

Four high-speed cameras, configured as shown in Fig. 1(C) were used to perform particle tracking in the roughly 8 × 8 × 4 cm^3^ measurement volume. The cameras were calibrated using Tsai’s method (Tsai, 1987), in which camera parameters were estimated from images of a dual-plane calibration plate. A subsequent volume self-calibration (VSC) was performed using low concentration tracer data to further refine and optimize the camera parameters (Wieneke, 2008). Four diffused infrared (IR) LED panels were used to illuminate the particles via scattered light. Most tracer particles appeared to be six to nine pixels in diameter on the images. Their positions were triangulated and tracked over time using our in-house 3D Lagrangian particle tracking code, OpenLPT (Tan et al., 2020). Velocity information was then obtained by convolving the trajectories with the derivative of a Gaussian, using an appropriate filter length (Voth et al., 2002).

With the given design parameters, Tan et al. (2023) predicts that the flow should be homogeneous and isotropic at a distance 6*M*_*j*_ = 66 cm away from the jet array. Thus, to ensure the jets are well-mixed, velocity field data was taken approximately 125 cm downstream of the jet array in the center of the tank. Spanwise and vertical profiles of the mean velocities ⟨*u*_*i*_⟩ and fluctuations 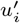 for one representative flow condition are shown in Fig. 2, with error bars indicating the standard deviation in the orthogonal directions. The mean velocity is approximately uniform in both directions, with the streamwise component ⟨*u*_1_⟩ deviating by only 3% on average from the centerline value, while the spanwise ⟨*u*_2_⟩ and vertical ⟨*u*_3_⟩ components remain near zero. The fluctuation components are similarly uniform and of comparable magnitude, consistent with approximately isotropic turbulence. Together, these results confirm that the jets are well-mixed within the measurement volume, as expected.

**Figure 2.**
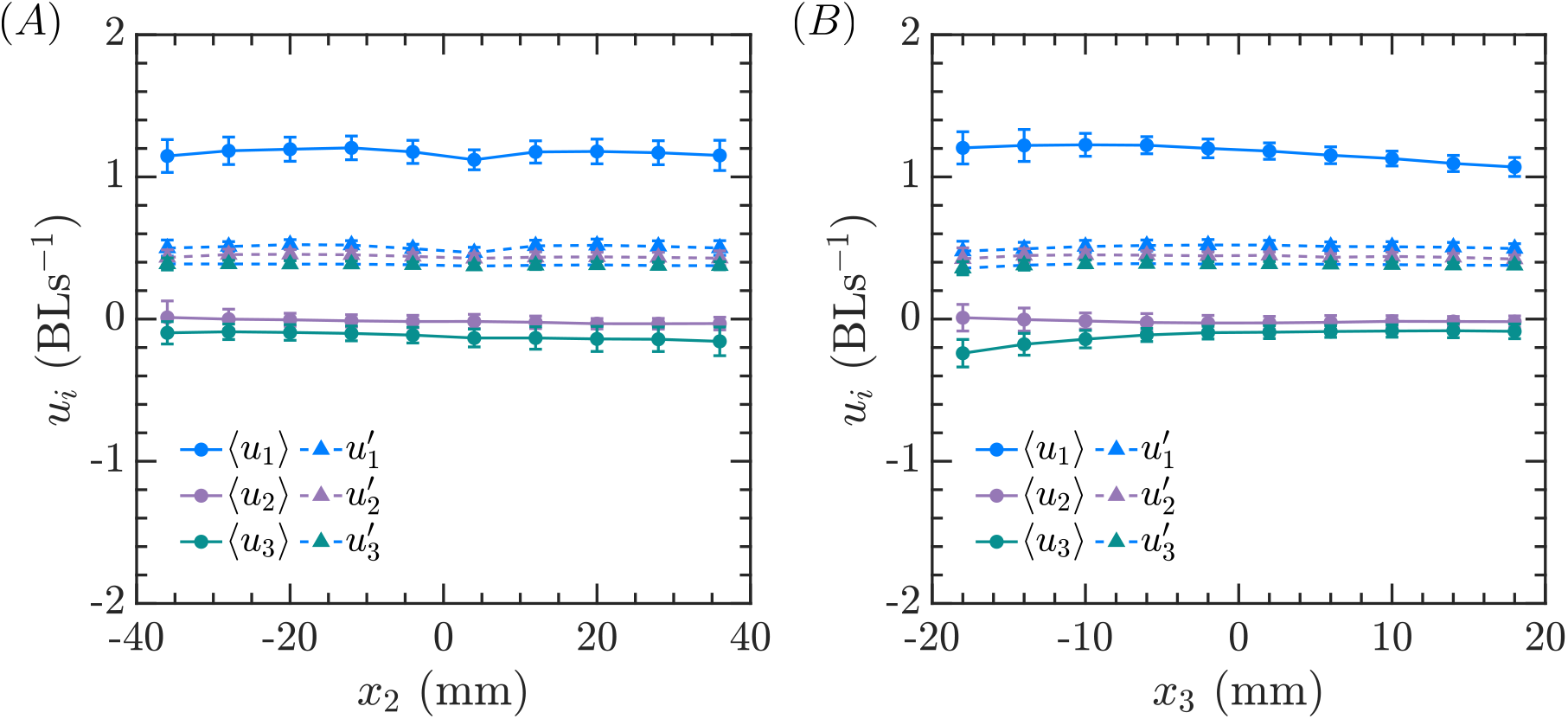
The mean and fluctuation velocity (⟨*u*_*i*_⟩ and 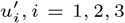 for three axes) are plotted as a function of (A) the spanwise axis (*x*_2_) and (B) the vertical axis (*x*_3_). The error bars indicate the standard deviation of the velocity in the orthogonal directions.

The fluctuation velocity 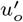 should scale linearly with the jet velocity *v*_*j*_ (Tan et al., 2023). And therefore, the turbulence intensity *I* should scale linearly with the injection ratio *J*, which is defined below, where *Q*_*j*_ is the total flow-rate of the jets, *Q*_*o*_ is the mean flow rate, *A*_*j*_ is the total cross-sectional area of the nozzles, *A*_*o*_ is the total cross-sectional area of the tank, and *ρ* is the density.

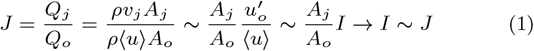

Fig. 3(A) shows the turbulence intensity generated by the jet array as a function of the injection ratio. Data points at an injection ratio of zero correspond to the baseline turbulence in the tank without the jet array installed. Velocities are normalized by the nominal body length of giant danio (*BL* = 5 cm). The results demonstrate that the linear relationship shown in Eq. 1 is observed for different combinations of mean flow rate and jet speed. Furthermore, it is shown that we can control the turbulence independent of the mean flow. For example, for a fixed swimming speed of 1 BLs^−1^, the turbulence intensity can be systematically varied from roughly 20% − 60%. In Fig. 3(B), it is shown that at this swimming speed the energy dissipation rate can also be systematically increased by increasing the injection ratio. The integral scale is roughly constant 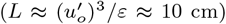 and the Kolomogorov scale *η* decreases with increasing energy dissipation rate. Thus, the jet array also provides a means for increasing the range of scales the fish will have to encounter. Here, the energy dissipation curves are determined from the 2nd (*D*_*LL*_, *D*_*NN*_) and 3rd (*D*_*LLL*_) order structure functions using Kolomogorov’s local isotropy hypothesis (Pope, 2000).

**Figure 3.**
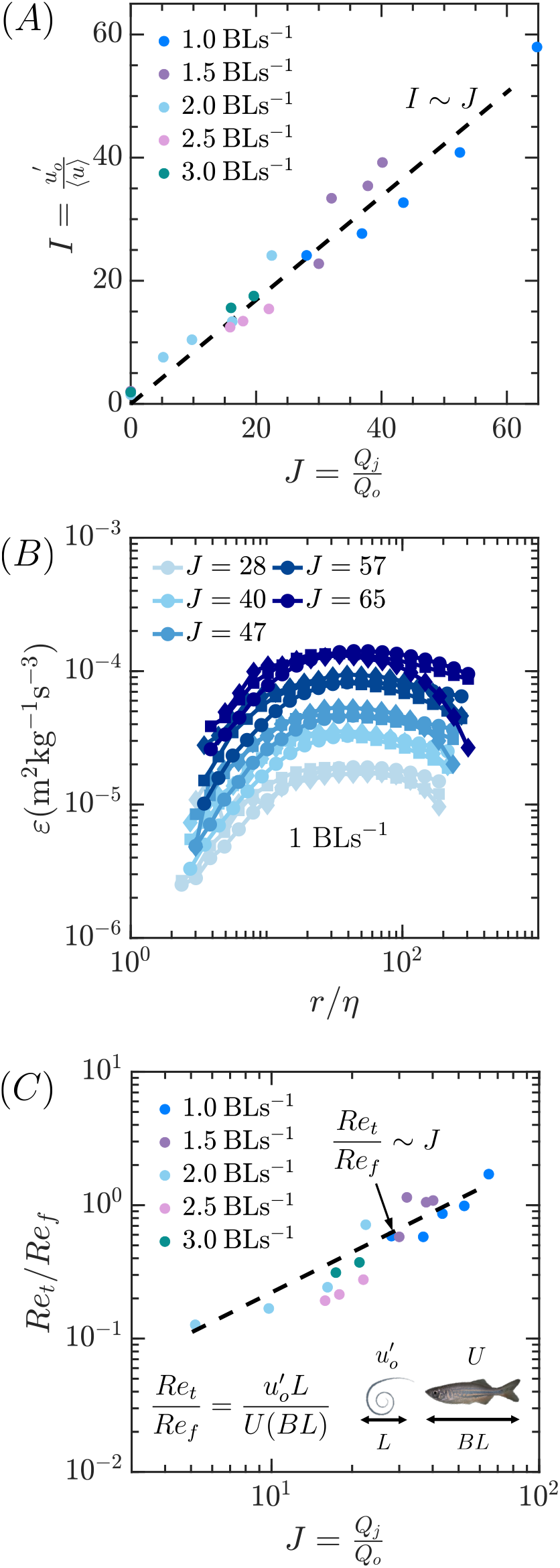
(A) Turbulent intensity *I* as a function of the injection ratio *J* demonstrating the expected linear scaling. (B) Energy dissipation rate (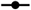 *D*_*LL*_, 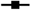 *D*_*NN*_,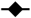 *D*_*LLL*_) at a fixed mean flow of 1 BLs^−1^ for different injection ratios demonstrating how the turbulence level can be systematically increased independently from the mean flow. (C) Ratio of the Reynolds number of the turbulence *Re*_*t*_ to that of the swimming fish *Re*_*t*_ which scales linearly with the injection ratio and spans an entire order magnitude.

Next, we demonstrate that the jet array enables systematic control of the relative scale of turbulence experienced by the fish. To quantify this, we compare the Reynolds number of the turbulence *Re*_*t*_ with that of the swimming fish *Re*_*f*_ . This ratio effectively compares the inertial forcing associated with the largest turbulent eddies to the characteristic inertial scale of the fish body.

Using the scaling established in Eq. (1), the ratio is expected to vary linearly with the injection ratio.

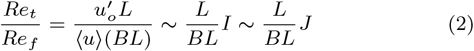

where *L* denotes the turbulence integral length scale and *BL* the fish body length. Notably, the viscosity cancels, so this ratio depends only on the relative velocity and length scales of the turbulence and the fish.

The predicted linear scaling is confirmed in Fig. 3(C), which shows that the jet array allows *Re*_*t*_*/Re*_*f*_ to be varied over approximately one order of magnitude (*Re*_*t*_*/Re*_*f*_ ≈ 0.1–1). For small *Re*_*t*_*/Re*_*f*_ values, turbulent fluctuations are weak relative to the fish’s inertial scale and are expected to have minimal influence on swimming behavior. As *Re*_*t*_*/Re*_*f*_ approaches unity, eddy-induced forcing becomes comparable to the fish’s inertia, and stronger hydrodynamic and behavioral effects are anticipated. Future work will quantify how these regimes actually influence locomotor performance.

A key advantage of the jet array developed here is its ability to systematically vary the turbulence characteristics through the controlled adjustment of various parameters. For example, increasing the injection ratio has been shown to increase the turbulence intensity while maintaining a fixed integral length scale. In addition, because the integral length scale grows and the energy dissipation rate decays with downstream distance, moving upstream would enable one to also investigate how fish respond to smaller, yet more energetic eddies. Further control of the turbulence characteristics can be achieved by modifying the jet spacing and nozzle diameter, which is feasible due to its simple and cost-effective design. Together, these capabilities provide the ability to test how fish respond to eddies that not only differ in size but also energy content, providing a flexible framework for addressing open questions about how turbulence influences the locomotion and behavior of a wide range of fish species.

### Perspective on future experiments

With FATE, a new class of experiments can be performed in which controlled turbulent environments are directly coupled with quantitative measurements of fish locomotion and behavior. A conceptual framework for these experiments is illustrated in Fig. 4. Turbulence intensity can be independently varied while keeping the fish velocity and length scales constant. Long time series of fish kinematics and behavior across multiple individuals will then be recorded, sampling over many tail-beat cycles and eddy turnover times. This ensures that both the intermittent nature of turbulence and the full behavioral dynamics are captured, enabling statistically converged comparisons with laminar baseline conditions to determine when turbulence begins to measurably affect fish.

**Figure 4.**
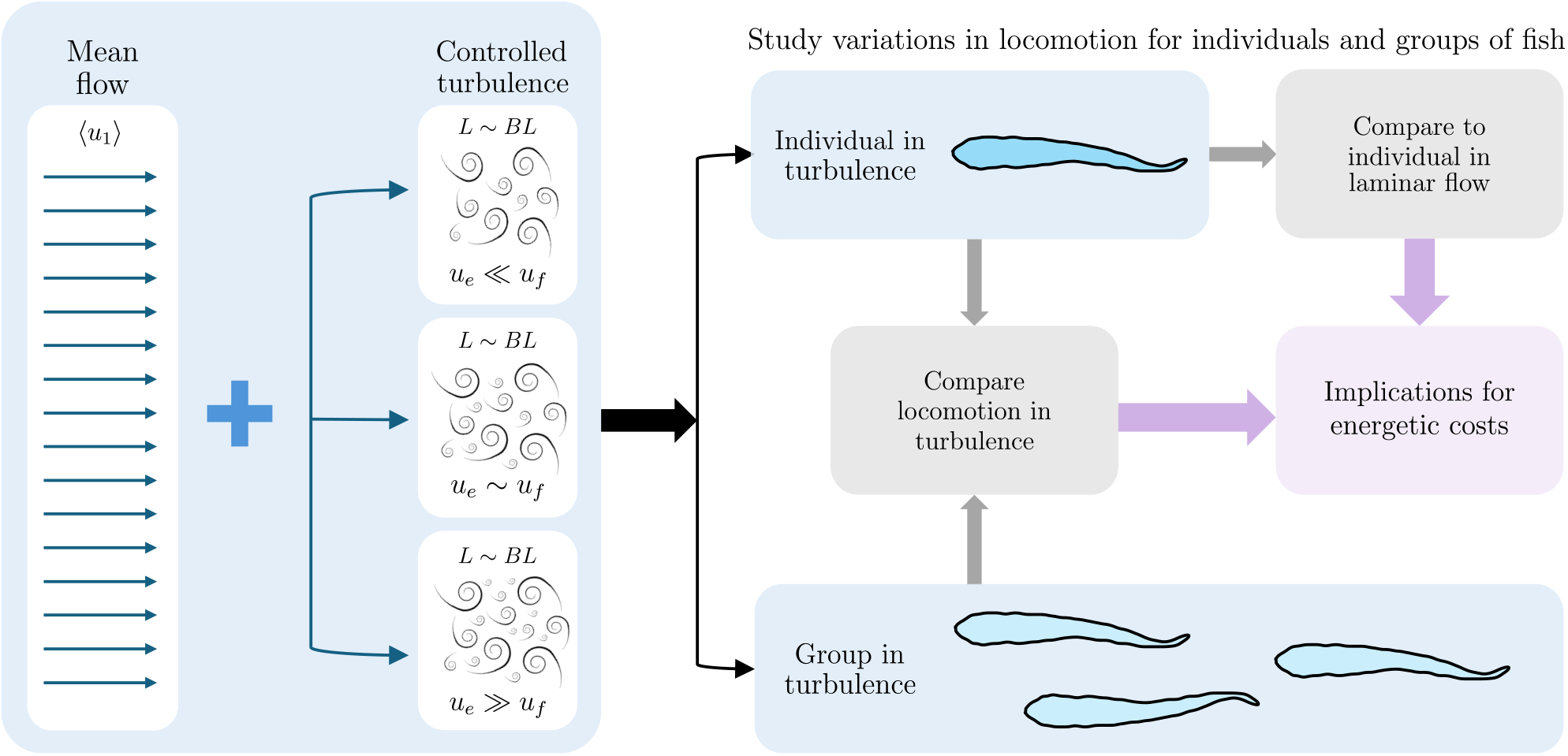
Conceptual framework for studying fish locomotion in turbulence. Turbulent flows should be generated such that specific quantities can be varied independently of the mean flow. For example, the integral length scale can be held constant while the eddy velocity scale *u*_*e*_ is systematically varied relative to the fish velocity scale *u*_*f*_ . For each condition, locomotion should be compared to that in laminar flows of matched mean velocity and to group locomotion in the same turbulent environment. Linking changes in kinematics and behavior to energetic costs is necessary to determine how turbulence affects performance and the extent to which collective behavior mitigates these effects

Once it is established when and how turbulence alters the swimming kinematics, the associated energetic consequences should then be quantified. Although predicting a fish’s energy expenditure is inherently challenging, prior studies suggest that the active metabolic rate scales with the tail beat kinematics (Sánchez-Rodríguez et al., 2023; Seo et al., 2025) since a fish’s undulatory motion generates the thrust force needed to overcome drag (Lighthill, 1969; Gazzola et al., 2014; Ventejou et al., 2025). This relationship provides a practical framework for estimating the additional energetic costs from measurable changes in swimming kinematics.

Consistent with this view, previous work reports that turbulence can increase tail beat amplitude *A* and frequency *f* (Pavlov and Skorobogatov, 2009; Zhang et al., 2024; Agbeti et al., 2024), promote more frequent fin deployment (McLaughlin and Noakes, 1998; Tritico and Cotel, 2010; Silva et al., 2012a; Maia et al., 2015; Webb, 2004; Duguay et al., 2018), and increase the occurrence of accelerations and corrective maneuvers (Maia et al., 2015; Silva et al., 2020; Li et al., 2022a). Such responses suggest fish are experiencing greater thrust demands in turbulence, which should correlate with larger energy expenditure.

For stationary swimmers, Pavlov and Skorobogatov (2009) has proposed the increased thrust demands comes from the increase in drag due to turbulence [⟨*D*⟩ = ⟨*u*⟩^2^(1 + *I*^2^)]. However, applying this formulation to the turbulence intensities reported in Zhang et al. (2024) substantially underpredicts the observed increase in swimming effort *Af* indicating that additional mechanisms likely contribute.

The proposed experiments will therefore look to directly quantify how modifications to the swimming gait, including changes in tail beat kinematics, fin deployment, and acceleration events, contribute to energy expenditure. In addition, they will determine whether these locomotor responses exhibit measurable signatures of the background turbulence, linking specific flow structures to distinct behavioral adjustments and increased energetic costs. Establishing these relationships will provide a more mechanistic and predictive framework for understanding how turbulent flows influence fish locomotion, behavior, and energy expenditure.

#### Role of collective behavior: energy savings

While identifying when and how turbulence may alter an individual fish’s locomotion and increase its energy expenditure is a necessary first step, it is equally important to understand the extent to which collective behavior can help fish mitigate these adverse effects. Schooling has been shown to reduce the metabolic rate, recovery time, and cortisol levels when compared to solitary swimmers in laminar flow conditions (Nadler et al., 2016; Zhang and Lauder, 2023; Schumann et al., 2023). Moreover, energy savings can be even more pronounced in turbulent flows, with reported maximum reductions of nearly 80% depending on the swimming speed (Zhang et al., 2024). To achieve these energy savings, fish within a school would be expected to exhibit less variation in their kinematics and perform fewer corrective maneuvers and accelerations compared to solitary fish in turbulence.

Testing this hypothesis requires extending the previous experiments to groups of fish and directly comparing their locomotion to that of solitary swimmers under identical flow conditions, as outlined in Fig. 4. Achieving this goal demands accurate, long-duration tracking of an individual’s kinematics within schools. Despite recent advances in multi-animal pose estimation and tracking, enabled by tools such as DeepLabCut (Lauer et al., 2022), SLEAP (Pereira et al., 2022), and TRex with DeepPoseKit (Graving et al., 2019; Walter and Couzin, 2021), persistent body overlap and occlusions continue to limit reliable extraction of individual motion, particularly when fish schools swim freely in three dimensions. Overcoming these limitations is therefore a prerequisite for quantifying how collective behavior alters locomotor effort in turbulence.

Even with improved tracking, kinematic measurements alone are insufficient to interpret reductions in effort; resolving the responsible hydrodynamic mechanisms is also required. One possible mechanism is that leading fish attenuate turbulence for downstream individuals while simultaneously generating more coherent, predictable signals through the shedding of vortex rings that trailing fish can exploit.

Confirming or refuting this hypothesis requires simultaneous measurements of the flow field and fish kinematics. Particle image velocimetry (PIV) can provide the necessary velocity information, aided by multi-angle illumination to reduce shadowing and reconstruction algorithms to fill sparse velocity regions (Gunes et al., 2006; Raben et al., 2012; Saini et al., 2016; Calicchia et al., 2023). Scanning laser systems further increase the likelihood of capturing multiple fish within the measurement plane and could even enable three-dimensional velocity and school reconstructions. A similar system has been utilized to study vertical migration of brine shrimp and can serve as a baseline for future set-ups (Fu et al., 2021; Mohebbi et al., 2024).

Both multi-angle PIV and scanning-laser systems can be integrated directly into FATE, making it a unique facility where turbulence can be systematically controlled and simultaneous fish–flow measurements obtained. Further advances in multi-animal tracking are still needed to reliably extract individual kinematics from dense, overlapping schools, but this experimental framework will enable more rigorous tests of how collective behavior reduces energy expenditure in turbulent flows.

#### Role of collective behavior: effective navigation

In addition to understanding the underlying mechanisms of collective behavior that mitigate the negative impacts of turbulence, it is also important to determine how the presence of conspecifics modulates the fish’s navigation strategy through heterogeneous flows. Due to the reduced energy expenditure, Goettel et al. (2015) suggested that western blacknose dace (*Rhinichthys obtusus*) may be more willing to traverse locally higher turbulent regions rather than expend energy searching for more favorable conditions. In contrast, Li et al. (2022a) observed that groups of Ya-fish (*Schizothorax prenanti*) increased their exploratory activity and were more effective at avoiding high mean, high turbulent regions than solitary swimmers. The strategy fish adopt may depend on the specific fish species, the flow conditions, and/or the group size, and thus warrants future investigation. However, identifying the mechanistic drivers that promote each strategy may be even more important.

This involves understanding how the collective behavior reduces the energy expenditure of the group, a topic discussed previously. It also requires determining how fish more efficiently locate favorable flow conditions when swimming in groups compared with swimming alone. Individual fish sense local flow information through the lateral line and integrate additional sensory modalities, such as vision, to guide navigation (Coombs and Janssen, 1989; Braun and Coombs, 2000; Montgomery et al., 2000; Bleckmann and Zelick, 2009; Ristroph et al., 2015). In the presence of conspecifics, however, motion is further influenced by social interaction forces. These forces are commonly categorized as attraction (maintaining cohesion), repulsion (avoiding collisions), and alignment (matching the direction of nearby neighbors) (Sumpter, 2006). The simultaneous influence of environmental sensing and social interactions creates a competition between responding to local flow cues and following the behavior of neighbors, which can modulate the schooling behavior (Filella et al., 2018; Ko et al., 2023; Peterson et al., 2024; Zhou et al., 2025a).

Berdahl et al. (2013) examined this interplay in golden shiners (*Notemigonus crysoleucas*), which have a preference for shaded habitats. These fish were tasked with navigating a noisy light environment to locate darker regions. They found that the efficiency with which darker regions were located increased with group size. Moreover, fish accelerations were more strongly correlated with the social vector (direction of social attraction) than with the environmental vector (direction of steepest ascent in the darkness level), suggesting that individuals prioritized social cues over direct environmental information. Similarly, Granger and Johnsen (2022) demonstrated that magnetoreceptive migrators, such as Fraser River sockeye salmon (*Oncorhynchus nerka*), can exploit collective navigation to improve migratory accuracy.

A heterogeneous turbulent flow field may similarly act as a noisy environment through which fish must navigate. Such a flow is generated in FATE by moving closer to the jet array (*x*_1_ *<* 6*M*_*j*_), where persistent turbulent jets create a repeating pattern of high mean, high turbulence regions. An example is shown in Fig. 5, which reports the mean velocity and turbulent kinetic energy (TKE) on a plane at the vertical mid-depth of the water, obtained from PIV. Giant danio positions from a representative snapshot are overlaid on the colormaps. Most fish locate the reduced velocity, reduced turbulence regions between jets. For the two isolated fish, the social vector points toward the group while the hydrodynamic vector points away from the jet region, indicating a conflict between social attraction and local flow avoidance. This conflict can influence not only individual movement decisions but also collective group dynamics. The resulting behavior likely depends on how unfavorable the intermediate swimming conditions are, as different flow regimes may elicit distinct responses. With the jet array, these intermediate conditions can be tuned by changing jet speed or nozzle diameter, which alters local mean velocities and turbulence scales and thus the severity of the flow patches fish must negotiate. Future work should quantify the relative influence of social interactions (the social vector) and local hydrodynamic cues (the hydrodynamic vector) on movement decisions to reveal how groups avoid highly turbulent regions while locating energetically favorable areas. FATE’s ability to independently tune local turbulence and mean flow provides the experimental control needed to disentangle these effects.

**Figure 5.**
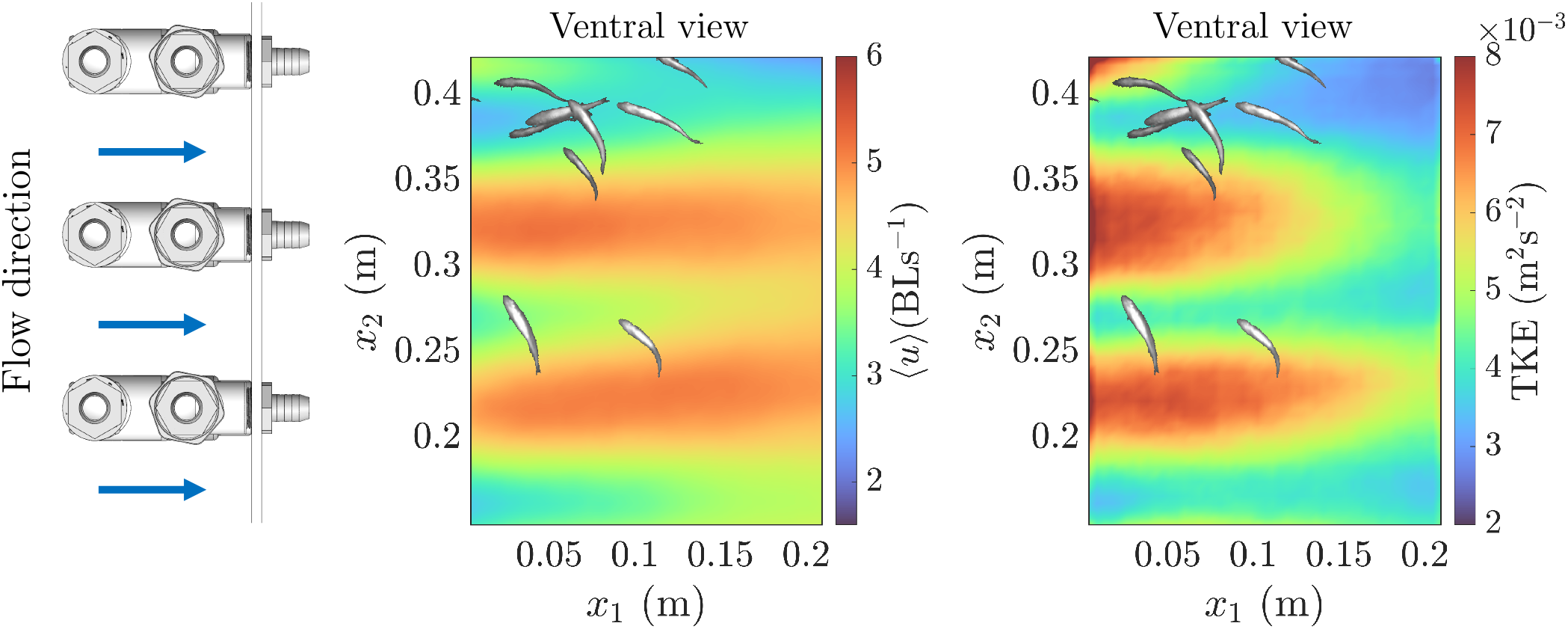
Representative slice of the mean velocity and turbulent kinetic energy approximately 30 cm downstream of the jet array, extracted at the vertical center of the channel, demonstrating the facility’s ability to generate challenging, heterogeneous flow conditions. Giant danio positions from a representative snapshot are overlaid on the colormaps. The majority of fish occupy a favorable low velocity, low turbulence region between jets, while isolated individuals must traverse high velocity, high turbulence regions to rejoin the group. The social vector drives movement toward the group, whereas the hydrodynamic vector promotes avoidance, creating competing cues that influence individual navigation and potentially affect collective behavior.

## Conclusion

It is well documented that fish tend to avoid flow conditions associated with elevated locomotor costs, such as turbulence. Although previous studies have begun to characterize how fish interact with the stochastic, multiscale nature of turbulent flows, many fundamental questions remain unresolved. In particular, two key areas warrant further investigation:

1. How does turbulence influence individual fish locomotion and behavior? Specifically, which turbulence characteristics drive responses, and how do they increase energetic costs?
2. What strategies do fish use to avoid or cope with turbulence? In particular, does collective behavior reduce energetic costs and help groups locate favorable flow regions, and what physical and behavioral mechanisms enable these benefits?

Addressing these questions will benefit from experimental approaches that decouple turbulence from the mean flow and enable independent control of the relevant spatial and temporal scales. We introduce a novel facility that achieves this control using a jet array, providing a framework to systematically vary turbulent scales relative to fish over an order of magnitude while simultaneously measuring behavioral responses. Future experiments will quantify statistically significant deviations in behavior and locomotion from baseline laminar conditions and apply scaling laws to assess the associated energetic costs. Tests with both solitary individuals and groups will help identify the physical and behavioral mechanisms that enable energy savings during collective swimming in turbulence. By shifting the measurement region closer to the jet array, heterogeneous flow fields can also be generated to examine how fish avoid or mitigate turbulence and navigate toward energetically favorable regions, both individually and as a group.

Understanding how fish truly interact with turbulence is motivated by both fundamental questions in animal locomotion and practical engineering needs. For example, turbulence has been shown to decrease passage efficiency and prolong migration times in fishways (Knapp et al., 2019). Insights from this line of research can therefore inform the design of these man-made structures, improving passage success while minimizing physiological stress and injury to fish. Moreover, autonomous underwater vehicles operate in similarly complex, unsteady flows. Thus, the principles uncovered here may also guide the development of robust, bio-inspired robotic systems capable of efficient navigation through turbulence.

## Animals and Housing

Giant danio (*Devario aequipinnatus*) with body length of approximately 5 cm and a height 1 cm. were procured from a local pet store (PetSmart). The fish were reared in multiple 76 L tanks. All tanks are equipped with temperature regulation (24-26 °C), aeration, and filtration systems. Water changes were performed weekly, and fish were fed commercial flake food daily. The animal holding and experimental procedures were approved by the Johns Hopkins Animal Care and Use Committee (IACUC) under protocol number FI24E79.

## Conflicts of interest

The authors declare that they have no competing interests.

## Funding

This work is supported by funds from the Office of Naval Research (N00014-21-1-2661 and N00014-24-1-2152).

## Data availability

The data underlying this article can be made available upon reasonable request.

## Author contributions statement

M.A.C and R.N. assisted in the design of the experimental facility and the conceptual framework for future work. M.A.C. aided in the assembly and construction of the facility, performed experiments, analyzed the data, wrote the manuscript, and made the visualizations. R.N. was responsible for funding acquisition.

## Acknowledgments

We thank Niel Leon, Rich Middlestadt, Mark Cooper, Daren Ayres, and Stipe Iveljic of WSE Manufacturing and Daniel Naugle, James Brady, Bob Klacka, Philip Craven, and other members of USA Composites for assisting in the design and manufacturing of the experimental facility. We also thank Xinyu Si, Bowen Zheng, and Shijie Zhong for their assistance in obtaining the 3D particle tracking data, and Ji Zhou for his feedback on the visualizations in this article.

## Notes

### Competing Interest Statement

The authors have declared no competing interest.

## References

M. V. Abrahams and P. W. Colgan. Risk of predation, hydrodynamic efficiency and their influence on school structure. Environmental Biology of Fishes, 13(3):195–202, 1985.

W. E. K. Agbeti, A. P. Palstra, S. Black, L. Magnoni, M. J. M. Lankheet, and H. Komen. Swimming at increasing speeds in steady and unsteady flows of atlantic salmon Salmo sapiens: Oxygen consumption, locomotory behaviour and overall dynamic body acceleration. Biology, 13(6):393, 2024.

I. Ashraf, H. Bradshaw, T.-T. Ha, J. Halloy, R. Godoy-Diana, and B. Thiria. Simple phalanx pattern leads to energy saving in cohesive fish schooling. Proceedings of the National Academy of Sciences of the United States of America, 114(37):9599– 9604, 2017.

J. B. Barlow, W. H. J. Rae, and A. Pope. Low-Speed Wind Tunnel Testing. John Wiley & Sons, New York, 3 edition, 1999.

D. N. Beal, F. S. Hover, M. S. Triantafyllou, J. C. Liao, and G. V. Lauder. Passive propulsion in vortex wakes. Journal of Fluid Mechanics, 549:385–402, 2006.

G. Bellani and E. A. Variano. Homogeneity and isotropy in a laboratory turbulent flow. Experiments in Fluids, 55(1):1646, 2014.

A. Berdahl, C. J. Torney, C. C. Ioannou, J. J. Faria, and I. D. Couzin. Emergent sensing of complex environments by mobile animal groups. Science, 339(6119):574–576, 2013.

H. Bleckmann and R. Zelick. Lateral line system of fish. Integrative Zoology, 4(1):13–25, 2009.

C. B. Braun and S. Coombs. The overlapping roles of the inner ear and lateral line: the active space of dipole source detection. Philosophical Transactions of the Royal Society of London. Series B: Biological Sciences, 355(1401):1115–1119, 2000.

E. Brown. Safe passage for fish: The case for in-stream turbines. Renewable and Sustainable Energy Reviews, 171:113074, 2023.

G. Căda, T. Carlson, J. Ferguson, M. Richmond, and M. Sale. Exploring the role of shear stress and severe turbulence in downstream fish passage. In Waterpower’99: Hydro’s Future: Technology, Markets, and Policy, pages 1–9. American Society of Civil Engineers, 1999.

M. A. Calicchia, R. Mittal, J.-H. Seo, and R. Ni. Reconstructing the pressure field around swimming fish using a physics-informed neural network. Journal of Experimental Biology, 226(8):jeb244983, 2023.

D. Carter, A. Petersen, O. Amili, and F. Coletti. Generating and controlling homogeneous air turbulence using random jet arrays. Experiments in Fluids, 57(12):189, 2016.

N. Cheng and A. W. Law. Measurements of turbulence generated by oscillating grid. Journal of Hydraulic Engineering, 127(3): 201–208, 2001.

S. Coombs and J. Janssen. Peripheral processing by the lateral line system of the mottled sculpin (cottus bairdi). In S. Coombs, P. Görner, and H. Münz, editors, The Mechanosensory Lateral Line, pages 299–319. Springer, New York, 1989.

A. J. Cotel and P. W. Webb. The challenge of understanding and quantifying fish responses to turbulence-dominated physical environments. In S. Childress, A. Hosoi, W. W. Schultz, and J. Wang, editors, Natural Locomotion in Fluids and on Surfaces, volume 155 of IMA Volumes in Mathematics and its Applications, pages 15–33. Springer, New York, NY, 2012.

A. J. Cotel and P. W. Webb. Living in a turbulent world—a new conceptual framework for the interactions of fish and eddies. Integrative and Comparative Biology, 55(4):662–672, Oct. 2015.

A. J. Cotel, P. W. Webb, and H. Tritico. Do brown trout choose locations with reduced turbulence? Transactions of the American Fisheries Society, 135(3):610–619, 2006.

M. B. Courtney, A. J. Flanigan, M. Hostetter, and A. C. Seitz. Characterizing sockeye salmon smolt interactions with a hydrokinetic turbine in the kvichak river, alaska. North American Journal of Fisheries Management, 42(4):1054– 1065, 2022.

M. Daghooghi and I. Borazjani. The hydrodynamic advantages of synchronized swimming in a rectangular pattern. Bioinspiration Biomimetics, 10(5):056018, 2015.

I. P. D. De Silva and H. J. S. Fernando. Oscillating grids as a source of nearly isotropic turbulence. Physics of Fluids, 6(7): 2455–2464, 1994.

V. Di Santo and E. Goerig. Swimming smarter, not harder: fishes exploit habitat heterogeneity to increase locomotor performance. Journal of Experimental Biology, 228 (Suppl1), 2025.

J. Duguay, B. Foster, R. W. Lacey, and T. R. Castro-Santos. Sediment infilling benefits rainbow trout passage in a baffled channel. Ecological Engineering, 125:38–49, 2018.

E. C. Enders, D. Boisclair, and A. G. Roy. The effect of turbulence on the cost of swimming for juvenile atlantic salmon (salmo salar). Canadian Journal of Fisheries and Aquatic Sciences, 60(9): 1149–1160, 2003.

A. Filella, F. Nadal, C. Sire, E. Kanso, and C. Eloy. Model of collective fish behavior with hydrodynamic interactions. Physical Review Letters, 120(19):198101, 2018.

M. K. Fu, I. A. Houghton, and J. O. Dabiri. A single-camera, 3d scanning velocimetry system for quantifying active particle aggregations. Experiments in Fluids, 62(8):168, 2021.

M. Gad-El-Hak and S. Corrsin. Measurements of the nearly isotropic turbulence behind a uniform jet grid. Journal of Fluid Mechanics, 62(1):115–143, 1974.

M. Gazzola, M. Argentina, and L. Mahadevan. Scaling macroscopic aquatic locomotion. Nature Physics, 10(10):758–761, 2014.

M. T. Goettel, J. F. Atkinson, and S. J. Bennett. Behavior of western blacknose dace in a turbulence modified flow field. Ecological Engineering, 58:414–422, 2015.

J. Granger and S. Johnsen. Collective movement as a solution to noisy navigation and its vulnerability to population loss. Proceedings of the Royal Society B: Biological Sciences, 289(1987):20221910, 2022.

J. M. Graving, D. Chae, H. Naik, L. Li, B. Koger, B. R. Costelloe, and I. D. Couzin. Deepposekit, a software toolkit for fast and robust animal pose estimation using deep learning. eLife, 8:e47994, 2019.

H. Gunes, S. Sirisup, and G. E. Karniadakis. Gappy data: To krig or not to krig? Journal of Computational Physics, 212(1):358–382, 2006.

S. F. Harding, R. P. Mueller, M. C. Richmond, P. Romero-Gomez, and A. H. Colotelo. Fish response to turbulence generated using multiple randomly actuated synthetic jet arrays. Water, 11(8): 1715, 2019.

T. E. Higham, W. J. Stewart, and P. C. Wainwright. Turbulence, temperature, and turbidity: The ecomechanics of predator–prey interactions in fishes. Integrative and Comparative Biology, 55 (1):6–20, July 2015.

S. G. Hinch and P. S. Rand. Swim speeds and energy use of upriver-migrating sockeye salmon (oncorhynchus nerka): role of local environment and fish characteristics. Canadian Journal of Fisheries and Aquatic Sciences, 55(8):1821–1831, 1998.

F. A. Hockley, C. A. M. E. Wilson, A. Brew, and J. Cable. Fish responses to flow velocity and turbulence in relation to size, sex and parasite load. Journal of the Royal Society Interface, 11(91): 20130814, 2014.

K. Ishikawa, H. Wu, S. Mitarai, and A. Genin. Turbulence effects on free and anchored zooplanktivorous fish in coral reefs. Coral Reefs, 2025. in press.

Y. Jiang, Z. Ma, and D. Zhang. Flow field perception based on the fish lateral line system. Bioinspiration Biomimetics, 14(4):041001, 2019.

J. L. Johansen, C. J. Fulton, and D. R. Bellwood. Avoiding the flow: refuges expand the swimming potential of coral reef fishes. Coral Reefs, 26(3):577–583, 2007.

M. Knapp, J. Montgomery, C. Whittaker, P. Franklin, C. Baker, and H. Friedrich. Fish passage hydrodynamics: insights into overcoming migration challenges for small-bodied fish. Journal of Ecohydraulics, 4(1):43–55, 2019.

H. Ko, G. Lauder, and R. Nagpal. The role of hydrodynamics in collective motions of fish schools and bioinspired underwater robots. Journal of the Royal Society Interface, 20(207):20230357, 2023.

M. K. Koiri, A. K. Sharma, A. Jha, and J. Kumar. A comprehensive review of bio-inspired swimming in robotic fishes. Sensors and Actuators A: Physical, 116913, 2025.

R. W. J. Lacey, V. S. Neary, J. C. Liao, E. C. Enders, and H. M. Tritico. The ipos framework: Linking fish swimming performance in altered flows from laboratory experiments to rivers. River Research and Applications, 28(4):429–443, Sept. 2011.

J. Lauer, M. Zhou, S. Ye, W. Menegas, S. Schneider, T. Nath, M. M. Rahman, V. Di Santo, D. Soberanes, G. Feng, V. N. Murthy, G. Lauder, C. Dulac, M. Weygandt Mathis, and A. Mathis. Multi-animal pose estimation, identification and tracking with deeplabcut. Nature Methods, 19(4):496–504, 2022.

G. Li, D. Kolomenskiy, H. Liu, B. Thiria, and R. Godoy-Diana. On the energetics and stability of a minimal fish school. PLOS ONE, 14(8):e0215265, 2019.

L. Li, M. Nagy, J. M. Graving, J. Bak-Coleman, G. Xie, and I. D. Couzin. Vortex phase matching as a strategy for schooling in robots and in fish. Nature Communications, 11(1):5408, 2020.

M. Li, R. An, M. Chen, and J. Li. Evaluation of volitional swimming behavior of Schizothorax prenanti using an open-channel flume with spatially heterogeneous turbulent flow. Animals, 12(6):752, 2022a.

M. Li, M. Chen, W. Wu, J. Li, and R. An. Differences in the natural swimming behavior of schizothorax prenanti individual and schooling in spatially heterogeneous turbulent flows. Animals, 13(6):1025, 2023.

Y. Li, Y. Xu, Z. Wu, L. Ma, M. Guo, Z. Li, and Y. Li. A comprehensive review on fish-inspired robots. International Journal of Advanced Robotic Systems, 19(4):172988062211037, 2022b.

J. C. Liao. A review of fish swimming mechanics and behaviour in altered flows. Philosophical Transactions of the Royal Society B: Biological Sciences, 362(1487):1973–1993, May 2007.

J. C. Liao and O. Akanyeti. Fish swimming in a k’arm’an vortex street: Kinematics, sensory biology and energetics. Marine Technology Society Journal, 51(5):48–55, 2017.

J. C. Liao, D. N. Beal, G. V. Lauder, and M. S. Triantafyllou. The k’arm’an gait: novel body kinematics of rainbow trout swimming in a vortex street. Journal of Experimental Biology, 206(6):1059– 1073, 2003a.

J. C. Liao, D. N. Beal, G. V. Lauder, and M. S. Triantafyllou. Fish exploiting vortices decrease muscle activity. Science, 302(5650): 1566–1569, 2003b.

M. J. Lighthill. Note on the hydromechanics of aquatic animal propulsion. Journal of Fluid Mechanics, 44(2):265–301, 1969.

H. Liu, J. Lin, D. Wang, J. Huang, H. Jiang, D. Zhang, Q. Peng, and J. Yang. Experimental study of the behavioral response of fish to changes in hydrodynamic indicators in a near-natural environment. Ecological Indicators, 154:110813, Oct. 2023.

M. A. Loehrke and H. M. Nagib. Control of free-stream turbulence by means of honeycombs: A balance between suppression and generation. Journal of Fluids Engineering, 98(3):342–351, 1976.

A. I. Lupandin. The effects of flow turbulence on the swimming speed of fish. Biology Bulletin, 32(5):461–466, 2005.

M. M. Macias, R. C. F. Mendes, T. F. Oliveira, and A. C. P. Brasil Junior. Hydrokinetic turbine impact assessment on fish. In L. Chen, editor, Advances in Clean Energy Systems and Technologies, Green Energy and Technology, pages 471–479. Springer, 2024.

A. Maia, A. P. Sheltzer, and E. D. Tytell. Streamwise vortices destabilize swimming bluegill sunfish (Lepomis macrochirus). Journal of Experimental Biology, 218(5):786–792, 2015.

S. Marras, S. S. Killen, J. Lindström, D. J. McKenzie, J. F. Steffensen, and P. Domenici. Fish swimming in schools save energy regardless of their spatial position. Behavioral Ecology and Sociobiology, 69: 219–226, 2015.

A. U. M. Masuk, A. Salibindla, S. Tan, and R. Ni. V-onset (vertical octagonal noncorrosive stirred energetic turbulence): A vertical water tunnel with a large energy dissipation rate to study bubble/droplet deformation and breakup in strong turbulence. Review of Scientific Instruments, 90(8):085105, Aug. 2019.

R. L. McLaughlin and D. L. Noakes. Going against the flow: an examination of the propulsive movements made by young brook trout in streams. Canadian Journal of Fisheries and Aquatic Sciences, 55(4):853–860, 1998.

R. D. Mehta and P. Bradshaw. Design rules for small low speed wind tunnels. Technical Report AGARD-AG-231, Advisory Group for Aerospace Research and Development (AGARD), NATO, Neuillysur-Seine, France, 1979.

A. Menzer, Y. Pan, G. V. Lauder, and H. Dong. Fish schools in a vertical diamond formation: Effect of vertical spacing on hydrodynamic interactions. Physical Review Fluids, 10(4):043104, 2025.

N. Mohebbi, J. Hwang, M. K. Fu, and J. O. Dabiri. Measurements and modelling of induced flow in collective vertical migration. Journal of Fluid Mechanics, 1001:A50, 2024.

J. Montgomery, G. Carton, R. Voigt, C. Baker, and C. Diebel. Sensory processing of water currents by fishes. Philosophical Transactions of the Royal Society of London. Series B: Biological Sciences, 355(1401):1325–1327, 2000.

S. Müller, V. Muhawenimana, G. Sonnino-Sorisio, C. A. M. E. Wilson, J. Cable, and P. Ouro. Fish response to the presence of hydrokinetic turbines as a sustainable energy solution. Scientific Reports, 13(1):7459, 2023.

L. E. Nadler, S. S. Killen, E. C. McClure, P. L. Munday, and M. I. McCormick. Shoaling reduces metabolic rate in a gregarious coral reef fish species. Journal of Experimental Biology, 219(18):2802– 2805, 2016.

D. A. Neitzel, D. D. Dauble, G. F. Cada, M. C. Richmond, G. R. Guensch, R. P. Mueller, C. S. Abernethy, and B. Amidan. Survival estimates for juvenile fish subjected to a laboratory-generated shear environment. Transactions of the American Fisheries Society, 133(2):447–454, 2004.

V. I. Nikora, J. Aberle, B. J. F. Biggs, I. G. Jowett, and J. R. E. Sykes. Effects of fish size, time-to-fatigue and turbulence on swimming performance: a case study of galaxias maculatus. Journal of Fish Biology, 63(6):1365–1382, Dec. 18 2003.

M. Odeh, J. F. Noreika, A. Haro, A. Maynard, T. R. Castro-Santos, and G.F. Čada. Evaluation of the effects of turbulence on the behaviour of migratory fish. Technical report (doe/bp-00000022-1), Bonneville Power Administration; U.S. Geological Survey, Portland, OR / Reston, VA, Mar. 1 2002.

C. S. Ogilvy and A. B. Dubois. The hydrodynamic drag of swimming bluefish (pomatomus saltatrix) in different intensities of turbulence: Variation with changes of buoyancy. Journal of Experimental Biology, 92(1):67–85, 06 1981. ISSN 0022-0949.

A. K. Othman, D. A. Zekry, V. Saro-Cortes, K. J. ee, and A. A. Wissa. Aerial and aquatic biological and bioinspired flow control strategies. Communications Engineering, 2(1), 2023.

Y. Pan, W. Zhang, J. Kelly, and H. Dong. Unraveling hydrodynamic interactions in fish schools: A three-dimensional computational study of in-line and side-by-side configurations. Physics of Fluids, 36(8):081909, 2024.

D. Pavlov and A. Kasumyan. Patterns and mechanisms of schooling behavior in fish: A review. Journal of Ichthyology, 40(Suppl. 2): S163–S231, 2000. ISSN 0032-9452.

D. Pavlov, A. Lupandin, and M. Skorobogatov. The effects of flow turbulence on the behavior and distribution of fish. J. Ichthyol., 40:S232–S261, 01 2000.

D. S. Pavlov and M. A. Skorobogatov. Effect of flow turbulence on the movement pattern of the caudal fin in fish. Doklady Biological Sciences, 428:464–466, 2009.

T. D. Pereira, N. Tabris, A. Matsliah, D. M. Turner, J. Li, S. Ravindranath, E. S. Papadoyannis, E. Normand, D. S. Deutsch, Z. Y. Wang, G. C. McKenzie-Smith, C. C. Mitelut, M. D. Castro, J. D’Uva, M. Kislin, D. H. Sanes, S. D. Kocher, A. L. Falkner, J. W. Shaevitz, and M. Murthy. Sleap: A deep learning system for multi-animal pose tracking. Nature Methods, 19(4):486–495, 2022.

A. N. Peterson, N. Swanson, and M. J. McHenry. Fish communicate with water flow to enhance a school’s social network. Journal of Experimental Biology, 227(17):jeb247507, 2024.

T. J. Pitcher, A. E. Magurran, and I. J. Winfield. Fish in larger shoals find food faster. Behavioral Ecology and Sociobiology, 10 (2):149–151, 1982.

S. B. Pope. Turbulent Flows. Cambridge University Press, Cambridge, UK, 2000. ISBN 9780521598866.

S. G. Raben, J. J. Charonko, and P. P. Vlachos. Adaptive gappy proper orthogonal decomposition for particle image velocimetry data reconstruction. Measurement Science and Technology, 23 (2):025303, 2012.

L. Ristroph, J. C. Liao, and J. Zhang. Lateral line layout correlates with the differential hydrodynamic pressure on swimming fish. Physical Review Letters, 114(1):018102, 2015.

D. G. Roche, M. K. Taylor, S. A. Binning, J. L. Johansen, P. Domenici, and J. F. Steffensen. Unsteady flow affects swimming energetics in a labriform fish (Cymatogaster aggregata). Journal of Experimental Biology, 217(3):414–422, 2014.

P. Saini, A. M. Steinberg, and C. M. Arndt. Development and evaluation of gappy-pod as a data reconstruction technique for noisy piv measurements in gas turbine combustors. Experiments in Fluids, 57(7):122, 2016.

J. Sánchez-Rodríguez, C. Raufaste, and M. Argentina. Scaling the tail beat frequency and swimming speed in underwater undulatory swimming. Nature Communications, 14(1):4136, 2023.

S. Schumann, G. Mozzi, E. Piva, A. Devigili, E. Negrato, A. Marion, D. Bertotto, and G. Santovito. Social buffering of oxidative stress and cortisol in an endemic cyprinid fish. Scientific Reports, 13 (1):20579, 2023.

J.-H. Seo, J. Zhou, and R. Mittal. Scaling laws for caudal fin swimmers incorporating hydrodynamics, kinematics, morphology, and scale effects. arXiv, abs/2507.11665, 2025.

A. T. Silva, J. M. Santos, M. T. Ferreira, A. N. Pinheiro, and C. Katopodis. Effects of water velocity and turbulence on the behaviour of iberian barbel (luciobarbus bocagei, steindachner 1864) in an experimental pool-type fishway. River Research and Applications, 27(3):360–373, 2011.

A. T. Silva, C. Katopodis, J. M. Santos, M. T. Ferreira, and A. N. Pinheiro. Cyprinid swimming behaviour in response to turbulent flow. Ecological Engineering, 44:314–328, 2012a. ISSN 0925-8574.

A. T. Silva, J. M. Santos, M. T. Ferreira, A. N. Pinheiro, and C. Katopodis. Passage efficiency of offset and straight orifices for upstream movements of iberian barbel in a pool-type fishway. River Research and Applications, 28(5):529–542, 2012b.

A. T. Silva, K. M. Bærum, R. D. Hedger, H. Baktoft, H.-P. Fjeldstad, K. Gjelland, F. Økland, and T. Forseth. The effects of hydrodynamics on the three-dimensional downstream migratory movement of atlantic salmon. Science of the Total Environment, 705:135773, 2020.

D. L. Smith, E. L. Brannon, and M. Odeh. Response of juvenile rainbow trout to turbulence produced by prismatoidal shapes. Transactions of the American Fisheries Society, 134(3):741–753, 2005.

D. L. Smith, E. L. Brannon, B. Shafii, and M. O. and. Use of the average and fluctuating velocity components for estimation of volitional rainbow trout density. Transactions of the American Fisheries Society, 135(2):431–441, 2006.

D. L. Smith, R. A. Goodwin, and J. M. Nestler. Relating turbulence and fish habitat: A new approach for management and research. Reviews in Fisheries Science Aquaculture, 22(2):123–130, 2014.

J. E. Stiansen and S. Sundby. Improved methods for generating and estimating turbulence in tanks suitable for fish larvae experiments. Scientia Marina, 65(2):151–167, 2001.

D. J. T. Sumpter. The principle of collective animal behavior. Philosophical Transactions of the Royal Society B: Biological Sciences, 361(1465):5–22, 2006.

J. C. Syms, M. A. Kirk, C. C. Caudill, and D. Tonina. A biologically based measure of turbulence intensity for predicting fish passage behaviours. Journal of Ecohydraulics, 9(3):55–67, 2021.

M. Taguchi and J. C. Liao. Rainbow trout consume less oxygen in turbulence: the energetics of swimming behaviors at different speeds. Journal of Experimental Biology, 214(9):1428–1436, 2011.

J. Tan, Z. Gao, H. Dai, Z. Yang, and X. Shi. Effects of turbulence and velocity on the movement behaviour of bighead carp (hypophthalmichthys nobilis) in an experimental vertical slot fishway. Ecological Engineering, 127:363–374, 2019.

S. Tan, A. Salibindla, A. U. M. Masuk, and R. Ni. Introducing openlpt: New method of removing ghost particles and high-concentration particle shadow tracking. Experiments in Fluids, 61(2):47, 2020.

S. Tan, X. Xu, Y. Qi, and R. Ni. Scalings and decay of homogeneous, nearly isotropic turbulence behind a jet array. Phys. Rev. Fluids, 8:024603, Feb 2023.

R. Thandiackal and G. V. Lauder. In-line swimming dynamics revealed by fish interacting with a robotic mechanism. eLife, 12: e81392, 2023.

G. Trinci, G. L. Harvey, A. J. Henshaw, W. Bertoldi, and F. Hölker. Life in turbulent flows: interactions between hydrodynamics and aquatic organisms in rivers. Wiley Interdisciplinary Reviews: Water, 4(3):e1213, 2017.

H. M. Tritico and A. J. Cotel. The effects of turbulent eddies on the stability and critical swimming speed of creek chub (semotilus atromaculatus). Journal of Experimental Biology, 213(13):2284–2293, 07 2010. ISSN 0022-0949.

R. Y. Tsai. A versatile camera calibration technique for high-accuracy 3d machine vision metrology using off-the-shelf tv cameras and lenses. IEEE Journal of Robotics and Automation, 3(4):323–344, 1987.

J. M. van der Hoop, M. L. Byron, K. Ozolina, D. L. Miller, J. L. Johansen, P. Domenici, and J. F. Steffensen. Turbulent flow reduces oxygen consumption in the labriform swimming shiner perch, Cymatogaster aggregata. Journal of Experimental Biology, 221(11):jeb168773, 2018.

B. Ventejou, T. Métivet, A. Dupont, and P. Peyla. Universal scaling laws for a generic swimmer model. Physical Review Letters, 134 (13):134002, 2025.

E. Villermaux, B. Sixou, and Y. Gagne. Intense vortical structures in grid-generated turbulence. Physics of Fluids, 7(8):2008–2013, 1995.

G. A. Voth, A. La Porta, A. M. Crawford, J. Alexander, and E. Bodenschatz. Measurement of particle accelerations in fully developed turbulence. Journal of Fluid Mechanics, 469:121–160, 2002.

T. Walter and I. D. Couzin. Trex, a fast multi-animal tracking system with markerless identification, and 2d estimation of posture and visual fields. eLife, 10:e64000, 2021.

P. W. Webb. Response latencies to postural disturbances in three species of teleostean fishes. Journal of Experimental Biology, 207 (6):955–961, 02 2004. ISSN 0022-0949.

P. W. Webb. Use of fine-scale current refuges by fishes in a temperate warm-water stream. Canadian Journal of Zoology, 84(8):1071– 1078, 2006.

P. W. Webb and A. J. Cotel. Waves and eddies: Effects on fish behavior and habitat distribution. In P. Domenici and B. G. Kapoor, editors, Fish Locomotion: An Eco-Ethological Perspective, pages 1–39. Science Publishers, Enfield, NH, 2010.

D. Weihs. Hydromechanics of fish schooling. Nature, 241(5387): 290–291, 1973.

B. Wieneke. Volume self-calibration for 3d particle image velocimetry. Experiments in Fluids, 45(4):549–556, 2008.

F. Yang and Y. Zeng. Collective swimming pattern and synchronization of fish pairs (gobiocypris rarus) in response to flow with different velocities. Journal of Fish Biology, 106(2): 442–452, 2025.

Y. Zhang and G. V. Lauder. Energy conservation by collective movement in schooling fish. eLife, 13:e90352, 2023.

Y. Zhang, H. Ko, M. A. Calicchia, R. Ni, and G. V. Lauder. Collective movement of schooling fish reduces the costs of locomotion in turbulent conditions. PLOS Biology, 22(6):e3002501, 2024.

J. Zhou, J. Seo, and R. Mittal. Effect of hydrodynamic wakes in dynamical models of large-scale fish schools. Physics of Fluids, 37(1):011912, 2025a.

J. Zhou, J.-H. Seo, and R. Mittal. Hydrodynamically beneficial school configurations in carangiform swimmers: Insights from a flow-physics informed model. Journal of Fluid Mechanics, 1014: A32, 2025b.

R. Zimmermann, H. Xu, Y. Gasteuil, M. Bourgoin, R. Volk, J. Pinton, E. Bodenschatz, and I. C. for Turbulence Research. The lagrangian exploration module: An apparatus for the study of statistically homogeneous and isotropic turbulence. Review of Scientific Instruments, 81(5):055112, 2010.

